# Activation of CP-AMPARs is required for homosynaptic and heterosynaptic structural LTP in the hippocampus

**DOI:** 10.1101/2025.10.16.682923

**Authors:** Laura A. Koek, Thomas M. Sanderson, Gregory Bond, John Georgiou, Benjamin Scholl, Graham L. Collingridge

## Abstract

Long-term potentiation (LTP) involves alterations in synaptic structure that are believed to underlie the persistent increase in synaptic efficacy. Here we compared structural LTP (sLTP) in EGFP-labelled spines with functional LTP, using field potential recording, at CA3-CA1 synapses in mouse hippocampal slices for ∼ 2 h following theta-burst stimulation (TBS). Activity-dependent labelling with FM4-64 allowed us to compare activated (FM+) and non-activated synapses and thereby compare homo- and hetero-synaptic sLTP. In addition, we related spine volume changes according to the probability of release, P(r), of activated synapses. At homosynaptic sites there was the expected NMDA receptor (NMDAR)-dependent potentiation of spine volume that persisted throughout the recording period. We found that this sLTP also required the synaptic activation of CP-AMPARs. There was also sLTP at heterosynaptic sites that, surprisingly, developed more quickly than the associated homosynaptic sLTP. This heterosynaptic sLTP was also dependent on the synaptic activation of both NMDARs and CP-AMPARs. Additionally, we observed a trans- and hetero-synaptic interaction, whereby the heterosynaptic spines grew according to the P(r) of the neighbouring active (homosynaptic) synapse. These observations have therefore advanced our understanding of sLTP in several ways; the first demonstration of the absolute requirement for the synaptic activation of CP-AMPARs for sLTP; the magnitude of heterosynaptic sLTP relative to homosynaptic sLTP and the hitherto unexpected combination of trans- and hetero-synaptic interactions.

## INTRODUCTION

Long-term potentiation (**LTP**) of synaptic transmission is known to involve the growth of dendritic spines^1–3^, a process that is believed to be important for learning and memory. Despite its relevance to cognition, the mechanisms that are responsible for structural LTP are only partially understood. For example, while it has been shown that spine growth involves the activation of NMDARs^1,2^, it is unclear whether other subtypes of ionotropic glutamate receptors are involved. In this regard, we and others have shown that the activation of calcium-permeable AMPARs (**CP-AMPARs**) are required for some forms of functional LTP^4–6^, although this is not invariably the case^7–9^ . We have shown further, using two-input experiments, that when CP-AMPARs trigger potentiation at synapses that receive high frequency stimulation (**HFS**), termed homosynaptic functional LTP, there is often a potentiation of nearby synapses that do not receive HFS, termed heterosynaptic functional LTP that also requires the activation of CP-AMPARs^9^. The purpose of the present experiments has therefore been to explore whether CP-AMPARs are involved in homosynaptic and heterosynaptic structural LTP (**sLTP**).

Most studies of sLTP have used uncaging of L-glutamate onto single spines^1,2,10–13^, a technique that allows for precise spatiotemporal control of plasticity induction and has led to considerable mechanistic knowledge. However, it differs from the physiological activation of spines that depends on the presynaptic stimulation rate and release probability, **P(r)**. Consequently, there is much less information regarding sLTP during physiologically-relevant, synaptically-triggered plasticity. A technical issue to overcome is that only a subset of synapses are activated by stimulation of the afferent fibres and so many imaged spines do not experience the direct effect of synaptic glutamate release. To overcome this issue, we used SynaptoRed C2 (FM4-64), an amphiphilic styryl dye that labels synaptic vesicle membranes during neurotransmitter release, to monitor activated synapses and we combined this method with two-photon structural dendritic spine imaging with concurrent field electrophysiology. This paradigm therefore allowed the study of both homosynaptic and heterosynaptic sLTP in response to synaptic induction.

Our findings show that the activation of CP-AMPARs is required for sLTP at both homosynaptic and heterosynaptic inputs. Interestingly, the level of heterosynaptic LTP is influenced by the P(r) of the nearby homosynaptic input. This cross-talk could enables stronger inputs to preferentially induce structural heterosynaptic potentiation that encodes associative learning and memory.

## RESULTS

### Methods for studying homosynaptic and heterosynaptic sLTP

Although traditionally thought of as an input-specific process^14^, there are many instances where functional LTP at the conditioned input induces functional LTP at non-conditioned inputs. Whether this heterosynaptic plasticity involves structural changes in spines was previously unknown. We therefore developed a method to study sLTP at both homosynaptic and heterosynaptic inputs at the same time (Fig. 1) in acutely-prepared hippocampal slices (Fig. 1A). We adapted methods that use FM dyes in dissociated cultures^15^ and organotypic hippocampal cultures^16^, as well as approaches to reduce background staining^17,18^, so that we could image both FM4-64 labelled (FM+) and unlabeled (FM−) boutons together with EGFP-expressing spines within hippocampal slices (Fig. 1B,D). The FM method is sufficiently sensitive that FM− boutons are either silent or have an extremely low P(r) of <0.1. Therefore, we combined field electrophysiology with simultaneous two photon imaging of EGFP spines and FM labelling (Fig. 1A-D) to distinguish homosynaptic and heterosynaptic sLTP, as well as probe for any relationships between sLTP and the P(r) at individual synapses. Following the FM staining and destaining protocol^16^ we made field potential recordings so that we could (i) ensure a stable baseline, of at least 30 min, before conditioning and (ii) monitor the level of LTP induced. For conditioning, we delivered a spaced TBS (sTBS) since this induces both homosynaptic and heterosynaptic LTP^5,6,19^. Due to fluorophore bleaching we limited the imaging to two baseline periods and 5 post-conditioning periods, spread over a duration of 180 min.

**Figure 1.**
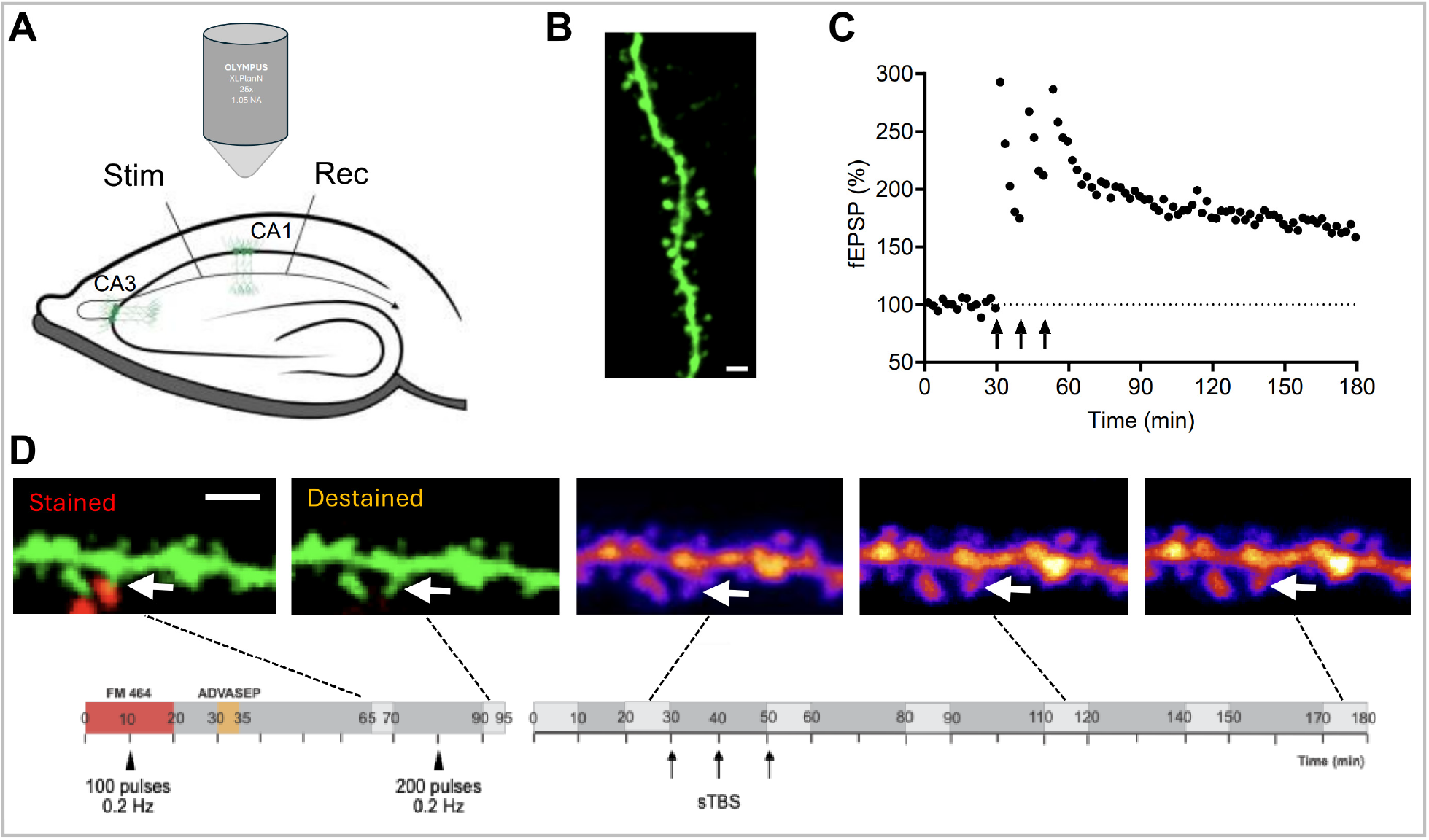
Measuring presynaptic release probability at activated CA1 dendritic spines. **(A)** Schematic of the experimental set-up consisting of multiphoton imaging with simultaneous electrophysiological field recordings at CA3–CA1 synapses within an *ex vivo* hippocampal slice from a Thy1-EGFP mouse. **(B)** Example of a dendritic branch from an EGFP+ CA1 neuron. Scale bar, 10 µm. **(C)** Example of functional LTP (normalised fEPSPs) time course that was induced (starting at time = 30 min, arrow) using a sTBS protocol (three rounds, every 10 min). **(D)** Experimental timeline (lower left) outlining the FM staining and destaining (coloured boxes), along with the subsequent (lower right) synaptic plasticity protocol (see STAR methods). Lighter gray boxes denote imaging sessions. Example images (upper left) of a dendrite (green) after FM staining (superimposed in red) and destaining (first 2 images), along with corresponding images (upper right) selected from one time point before and two time points after the sTBS delivery to induce structural LTP (sLTP).

### Structural LTP requires the synaptic activation of CP-AMPARs

Delivery of a sTBS protocol resulted in both functional LTP (Fig. 2A) and sLTP (Fig. 2B). Consistent with numerous previous studies, the functional LTP comprised an initial short-term potentiation (STP) followed by a fairly stable functional LTP (Fig. 2A). Quantified 2 h following the final TBS, the field excitatory postsynaptic potential (fEPSP) increased to 157 ± 6 % (values expressed as the mean ± SEM throughout; n=12). When all spines (both FM+ and FM−) were analyzed together there was sLTP that persisted for the duration of the recording (∼2 h). However, there was no structural correlate of STP. The sLTP was evident by 10 min following the TBS (the earliest time point measured) and remained stable throughout. Quantified from 120 min to 180 min, the spine volume increased to 120 ± 5 % (n = 186 spines).

**Figure 2.**
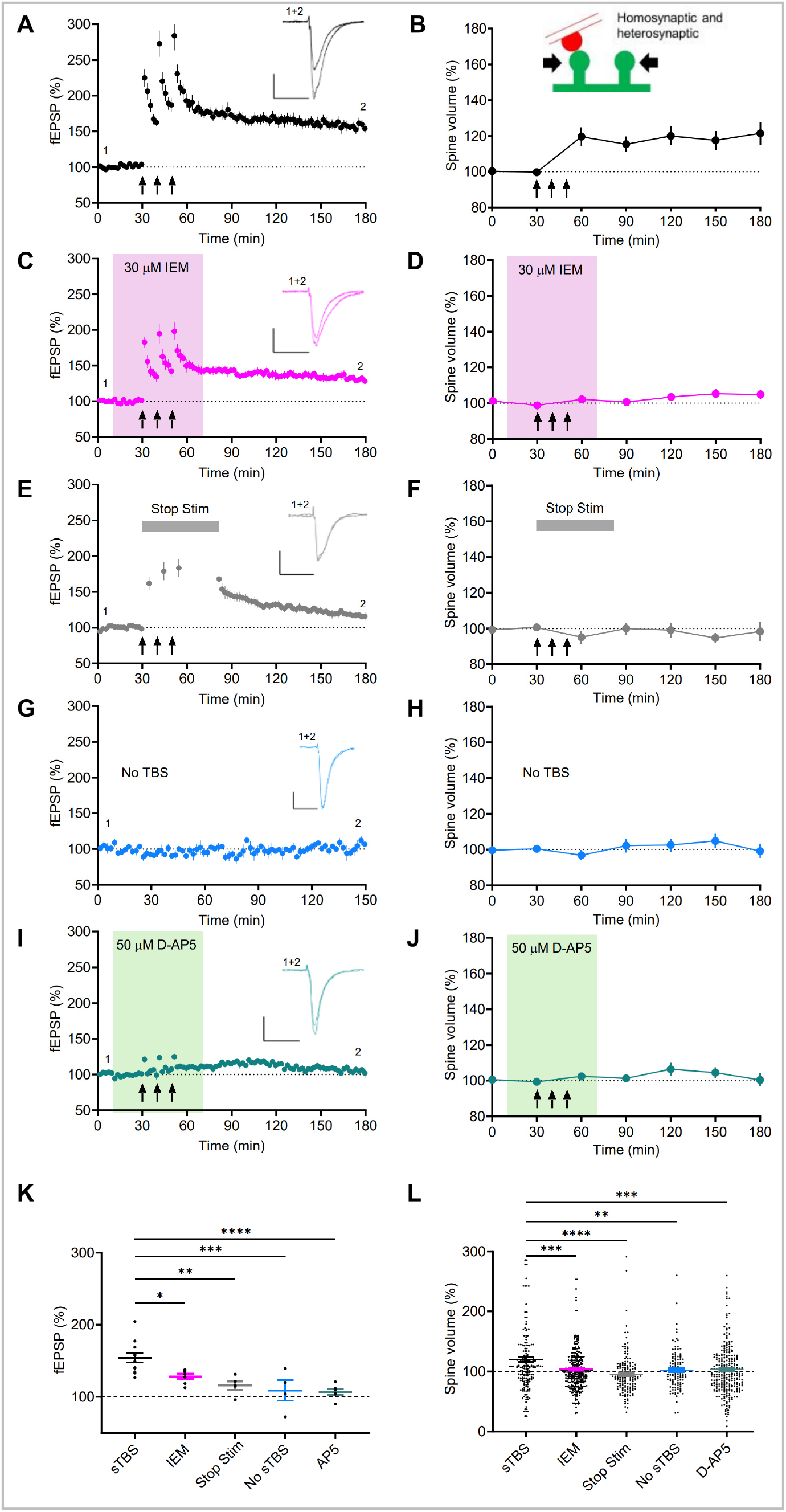
sLTP requires the synaptic activation of CP-AMPARs. **(A)** Time-course plot showing average LTP obtained in control conditions. **(B)** Time-course plot showing average sLTP obtained. Both activated (homosynaptic, FM+) and adjacent heterosynaptic spines (FM−) within 10 µm were included in this analysis. **(C)** Time-course plots showing less LTP was obtained in the presence of a CP-AMPAR inhibitor (IEM). **(D)** No corresponding sLTP in the presence of IEM. **(E)** Time-course plot showing lack of LTP when baseline stimulation is stopped (gray bar). **(F)** Corresponding time course plot of sLTP, showing lack of sLTP. **(G)** Time-course plot showing stable baseline when no TBS is delivered. **(H)** Tme-course plot showing stable spine volume when no TBS is delivered. **(I)** LTP is blocked in the presence of the NMDAR inhibitor D-AP5. **(J)** corresponding time course plot showing lack of sLTP in the presence of AP5. **(K)** Summary of individual data showing mean (± s.e.m) level of synaptic responses achieved (LTP) at the end of the experiments. **(L)** Individual and mean (± s.e.m) corresponding sLTP from all of the imaging data. The fEPSP traces are the average of five consecutive responses obtained from a representative experiment at the times indicated by labels ‘1’ and ‘2’. Calibration bars: 1 mV / 5 ms. *p<0.05, ** p<0.01, ***p<0.001 (see text for precise p-values).

Treatment with IEM-1460 (IEM), at a concentration that selectively affects CP-AMPARs at these synapses^6,9,20,21^ reduced, but did not abolish, functional LTP (Fig. 2C), which is consistent with our previous findings. The level of functional LTP induced at 120 min post conditioning was 133 ± 3% (n = 7), which was significantly less than that observed under control conditions (p = 0.007). In contrast, IEM completely prevented the induction of sLTP (Fig. 2D). As a second test for the involvement of CP-AMPARs, we stopped stimulation (**Stop Stim**) after each TBS episode (apart from 4 stimuli to monitor the level of STP) and for 30 min post TBS (Fig. 2E). This protocol also selectively prevents the component of LTP that requires the synaptic activation of CP-AMPARs^22^. With Stop Stim, the level of LTP induced at 120 min post conditioning was 116 ± 4 % (n=5), which was again significantly different to that observed under control conditions (p < 0.0001). As before, despite significant functional LTP, sLTP was eliminated (Fig. 2F).

As a control for recording stability, we monitored baseline stimulation without delivering sTBS. This resulted in neither any functional (Fig. 1G) nor structural (Fig. 1H) changes from baseline. As a second control we blocked NMDA receptors (**NMDARs**) by applying D-AP5, which prevents both functional^23^ and structural LTP^24^. Again, there were no changes in in either the functional (Fig. 1I) or structural (Fig. 1J) responses, demonstrating both processes are NMDAR-dependent. The level of functional LTP for each individual slice recording is plotted in Fig. 2K and the level of structural LTP for each spine measurement is shown in Figure 2L. We can also conclude, that whereas functional sTBS-induced LTP is partially dependent on the synaptic activation of CP-AMPARs, structural LTP is entirely dependent on their activation.

### Homosynaptic and heterosynaptic LTP both require the synaptic activation of CP-AMPARs

We next subdivided the spines into those adjacent to FM+ and FM− boutons (Fig. 3). As expected, the FM+ spines displayed sLTP (Fig. 3A); the average level of sLTP was 123 ± 9 % (n = 50 spines) over the 120 min to 180 min timepoints. Interestingly, there was a tendency for the spine volume to increase gradually from 90 min, reaching a maximum at 150 min of 131 ± 12 % (n = 41). Somewhat surprisingly, the FM− spines showed a quicker rate of increase in sLTP, (Fig. 3B); but reached a similar level of final sLTP averaged over the 120 to 180 min timepoints of 119 ± 5 % (n = 136). Consistent with the combined analysis of FM+ and FM− boutons (Fig. 2), IEM treatment prevented sLTP at both FM+ (Fig. 3C) and FM− (Fig. 3D) synapses.

**Figure 3.**
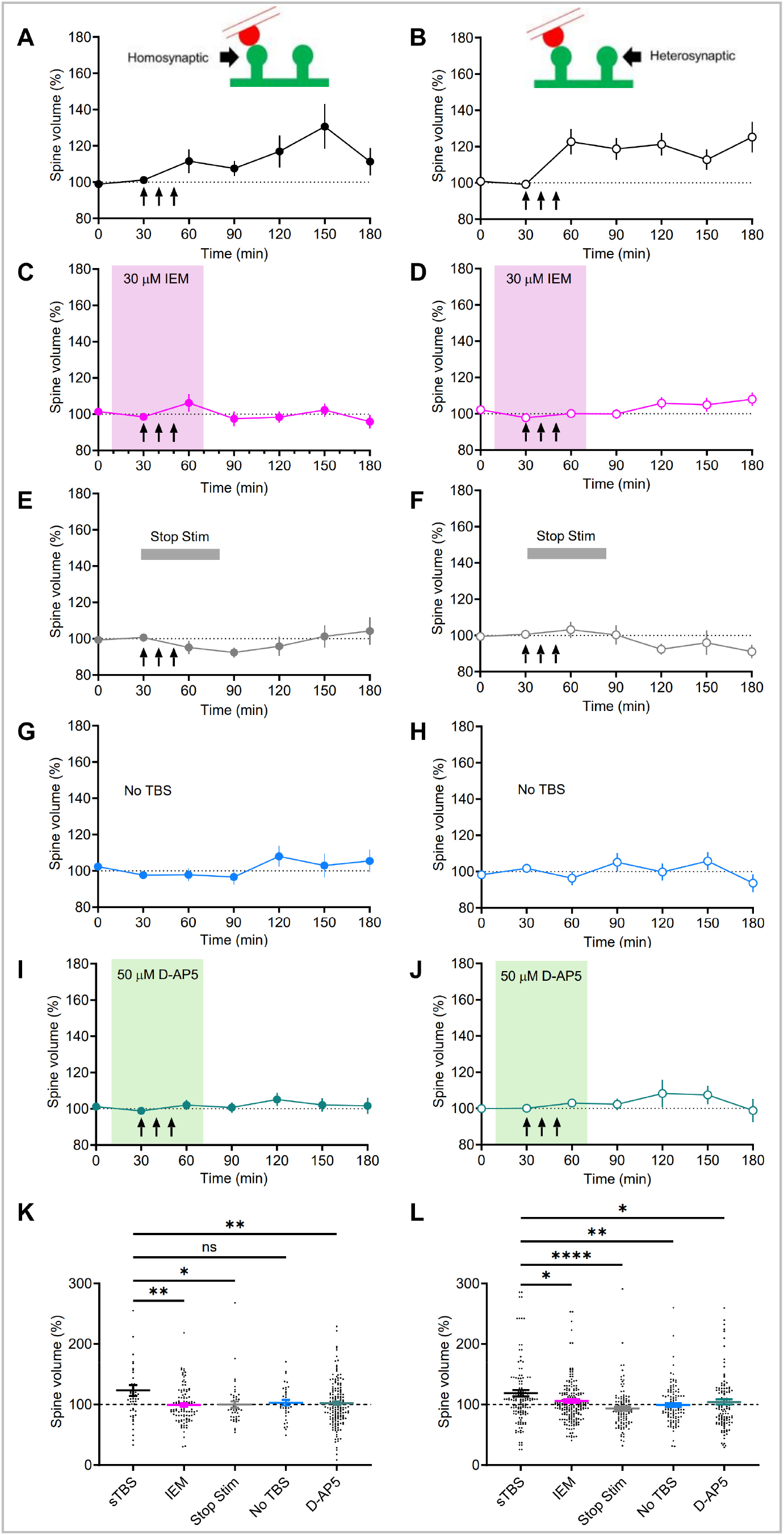
Both homosynaptic and heterosynaptic sLTP depend on CP-AMPAR insertion. **(A)** Time-course plot showing sLTP for only the homosynaptic spines following sTBS induction. **(B)** Time-course plot showing sLTP for the heterosynaptic spines. **(C)** Time-course plots showing a lack of homosynaptic sLTP when sTBS was delivered in the presence of the CP-AMPAR inhibitor IEM. **(D)** Heterosynaptic sLTP is blocked in the presence of IEM. **(E)** Time-course plot showing lack of homosynaptic sLTP when baseline stimulation is stopped (gray bar). **(F)** No heterosynaptic sLTP during stop stimulation. **(G)** Time-course plot showing absence of any homosynaptic sLTP when only baseline stimulation is delivered without any TBS. **(H)** Time-course plot showing heterosynaptic spine volume is unchanged when no TBS is delivered. **(I)** Homosynaptic sLTP is blocked in the presence of NMDAR inhibitor D-AP5. **(J)** Heterosynaptic sLTP is also blocked in the presence of D-AP5. **(K)** Individual spine data with mean (± s.e.m) level of homosynaptic sLTP in each type of experiment. **(L)** Mean (± s.e.m) of heterosynaptic sLTP.

Similarly, pausing baseline stimulation prevented sLTP at both FM+ (Fig. 3E) and FM− (Fig. 3F) synapses. Baseline stimulation alone had no effect at either FM+ (Fig. 3G) or FM− (Fig. 3H) spines. Similarly, conditioning in the presence of D-AP5 had no effect on either FM+ (Fig. 3I) or FM− (Fig. 3J) synapses. Analysis of individual FM+ and FM− synapses are illustrated in Figures 3K and L, respectively.

### Heterosynaptic plasticity is affected by the initial P(r) of the conditioned synapse

Next, we determined whether structural homosynaptic LTP was affected by P(r) of the synapse. Figure 4B shows examples of low P(r) and high P(r) synapses and Figure 4C the P(r) distribution for 622 synapses. We arbitrarily divided synapses into low P(r) and high P(r) based on whether their P(r) was below or above 0.2, respectively. There was no significant difference spine volume growth between high and low P(r) synapses although we observed a tendency for low P(r) synapses to grow more slowly and for the effect to be more sustained (Fig. 4D).

**Figure 4.**
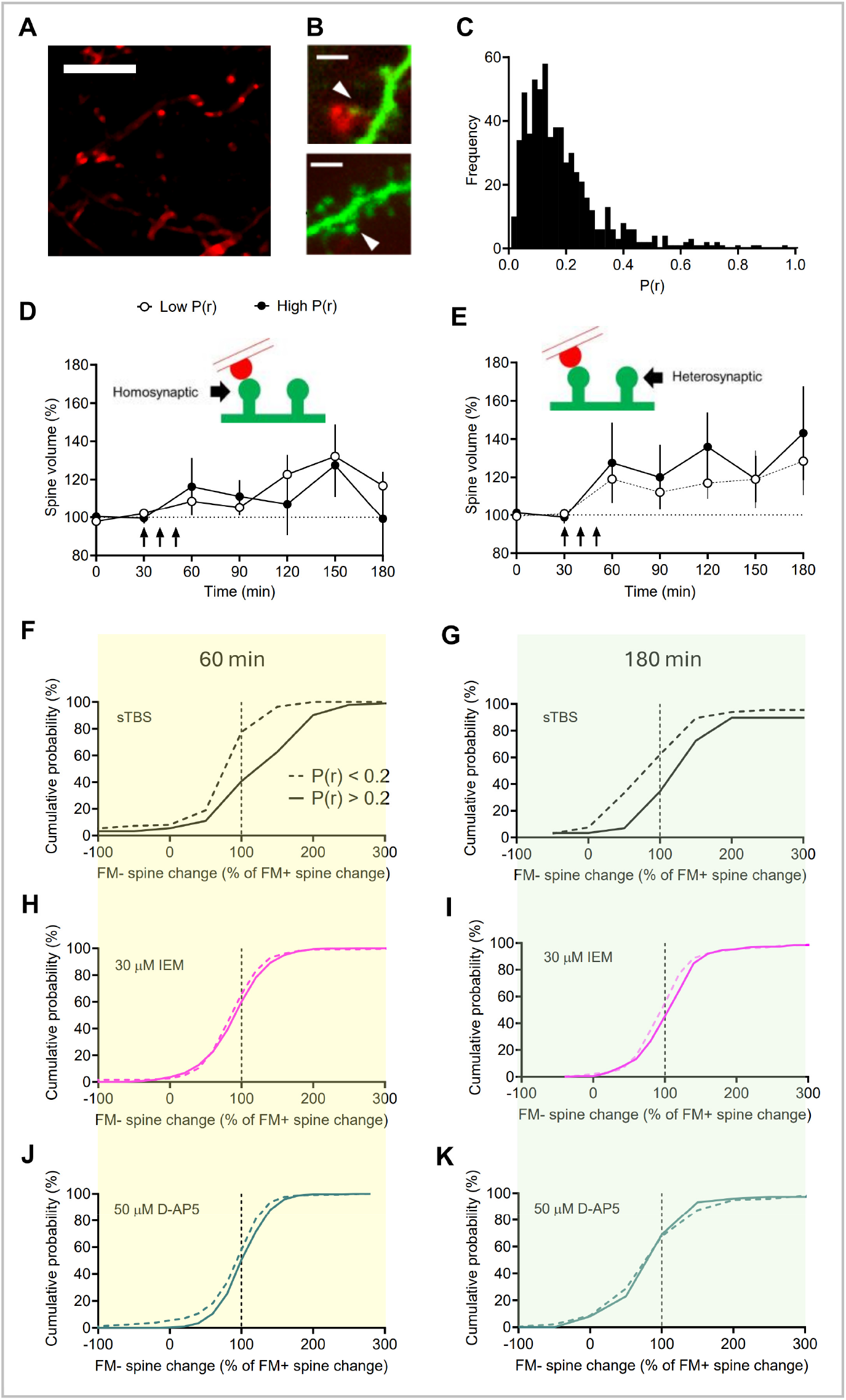
Heterosynaptic sLTP depends on CP-AMPARs and release probability of the activated synapse. **(A)** Image showing FM puncta (red). Scale bar, 10 µm. (B) Representative images showing homosynaptic EGFP+ (green) spines (white arrowheads) receiving inputs that have either a high P(r) [P(r) = 0.48; upper image] or low P(r) [P(r)= 0.16; lower image]. Scale bar, 10 µm. **(C)** Histogram of P(r) distribution based on FM fluorescence intensity. **(D)** Time-course plots showing homosynaptic sLTP, split into low (open circles) and high P(r) (filled circles) under sTBS conditions. **(E)** Time-course plots of heterosynaptic sLTP, split into proximity (within 10 µm) to low (open circles) versus high P(r) (filled circles). **(F)** Cumulative distribution plots showing heterosynaptic spines proximal to high P(r) spines had higher relative growth than those proximal to low P(r) spines. p = 0.561 for early sLTP (60 min). **(G)** Statistically significant more growth (p = 0.0188) during late sLTP (180 min) for high P(r) heterosynaptic spines. **(H)** No heterosynaptic sLTP in the presence of IEM during early sLTP; **(I)** nor at late sLTP time-point. **(J)** No sLTP in the presence of D-AP5 at early time point after TBS; **(K)** nor at end of experiment.

We next determined whether structural heterosynaptic LTP was affected by P(r) of the neighbouring active synapse. There was no significant difference between high and low P(r) synapses (Fig. 4D) though there was a tendency for the level of sLTP to increase with time and for the spines close to high P(r) synapses to grow more (Fig. 4E). To examine this potential heterosynaptic relationship in more detail, we plotted the change in heterosynaptic spine (FM−) volume as a percentage of the sLTP of the nearest homosynaptic (FM+) spine. We did this for low P(r) (<0.2) and high P(r) (>0.2) synapses as before, and at 2 timepoints: immediately after sTBS at the 60 min timepoint (Fig. 4F) and at the end of the experiment at the 180 min timepoint (Fig. 4G). We observed that heterosynaptic spines close to homosynaptic synapses with high P(r) mainly grew to a larger degree than the activated spines themselves (exhibiting a rightwards shift above 100% sLTP of the homosynaptic FM+ synapse). Conversely, spines close to low P(r) synapses mainly decreased relative to the activated spine (Fig. 4F,G). This differential effect of P(r) was not observed when sTBS was delivered in the presence of IEM (Fig. 4H,I), or when TBS was delivered in the presence of D-AP5 (Fig. 4J,K).

These observations therefore demonstrate a relationship between structural heterosynaptic plasticity and the P(r) of neighbouring active synapses.

## DISCUSSION

In the present study we have studied structural and functional LTP induced by spaced TBS at hippocampal CA3–CA1 synapses and have revealed several new properties of structural LTP. First, we have found that sLTP requires the synaptic activation of CP-AMPARs. Second, that structural homosynaptic LTP is associated with structural heterosynaptic LTP that also requires the synaptic activation of CP-AMPARs. Third, the extent of heterosynaptic LTP is influenced by the P(r) of the activated synapse. These studies have therefore extended the understanding of the mechanistic basis of sLTP, a process that is important for learning and memory.

### Homosynaptic structural LTP

Although there have been many studies of sLTP induced by local uncaging of glutamate^1,2,10–13^, studies that examine structural LTP in response to electrical synaptic activation are scarce. Our study most closely resembles that of Yang et al^25^, who delivered TBS and studied structural LTP within a 20-30 µm radius of the stimulation site. Although there are several similarities between the two studies there are also differences that can be explained, at least in part, by differences in the protocols used. First, the functional LTP in our study developed rapidly in contrast to the more slowly developing LTP, taking ∼30 min to reach a maximum level, in the study by Yang et al^25^. This was most likely because in our study the initial potentiation was STP, a form of synaptic potentiation that decays in a strictly activity-dependent manner and is probably largely presynaptic in origin^26^. We suspect that if STP is subtracted then the LTP time-course in the two studies would be similar. Consistent with a gradually developing LTP, in earlier studies we observed that postsynaptic sensitivity of neurons to ionophoretically applied AMPAR ligands developed with a similar time course^27^. Many subsequent studies have also documented an LTP that develops gradually (e.g.,^28^). Our results contrast with studies using uncaging of glutamate, where the spine changes occur rapidly^29^ . The distinct structural growth profiles presumably relate to differences in the induction protocol. With uncaging, a single spine is activated rapidly whereas with synaptic stimulation a single spine is activated much more intermittently at a rate that depends on the rate of stimulation post TBS and the P(r) of the synapse and is possibly also influenced by the rate of spontaneous activity.

In terms of sLTP, in the study of Yang et al (2008)^25^ there was a rapid, step change increase in spine volume that showed no changes for the 45 min recording period. We limited the number of measurements that we made over this period so that we could follow structural LTP for longer. We found that sLTP persisted for at least 2 h post TBS. This time-course is consistent with the slow development of postsynaptic sensitivity to AMPAR ligands, in response to synaptically-induced LTP.

### Heterosynaptic sLTP

By using FM4-64, we could distinguish synapses that were activated during the axonal electrical stimulation and those that were not activated. Analysis of the fluorescence intensity distribution indicates that we could detect very low P(r) synapses, such that the FM− ones are truly silent (or of extremely low Pr). We suspect that the majority of these are synapses were not activated by the stimulating electrode due to the orientation of the axons relative to the slice preparation. Although heterosynaptic structural LTP has not previously been studied in response to axonal stimulation, the principle that this process can occur has been demonstrated using glutamate uncaging^13^.

Previously, we observed functional heterosynaptic LTP under conditions where CP-AMPARs were activated. The present finding that structural heterosynaptic LTP also requires activation of CP-AMPARs suggests that the two processes are related. Surprisingly, however, structural heterosynaptic LTP is of greater magnitude than functional heterosynaptic LTP, possibly reflecting a non-linearity between an increase in spine volume and the increase in the number of newly inserted AMPARs. With homosynaptic LTP, the mismatch in the opposite direction likely reflects the sizeable component of LTP that is independent of CP-AMPARs and may represent the insertion of new AMPARs synapses without the need for structural alterations.

### The influence of P(r) on sLTP

The observation that the P(r) of a synapse influences the likelihood that neighbouring FM− spines will grow, demonstrates a hitherto unexpected trans-synaptic interaction. Since this effect was abolished by IEM-1460, it seems most likely that high P(r) synapses result in greater postsynaptic activation of CP-AMPARs, which then mediate the heterosynaptic interactions.

### The mechanism of homosynaptic and heterosynaptic sLTP

The principal observation in the present study is that both homosynaptic and heterosynaptic structural LTP are eliminated by inhibition of CP-AMPARs. In contrast, and consistent with our previously reports^9,20,21^, functional LTP was reduced but not abolished when CP-AMPARs were blocked.

It is established that functional LTP comprises two mechanistically distinct forms of synaptic potentiation, based on sensitivity to inhibition of CP-AMPARs, PKA and *de novo* protein synthesis. We refer to the insensitive form as LTP1 (also known as early-LTP) and the sensitive component as LTP2 (also known as late LTP). The complete inhibition of sLTP by IEM and by pausing stimulation therefore suggests that structural LTP corresponds to LTP2. Consistent with this, sLTP induced by uncaging ^29^or TBS^6^ is also sensitive to inhibitors of PKA and protein synthesis.

We have proposed previously that heterosynaptic functional LTP is due to spread of Ca^2+^, via Ca^2+^-induced Ca^2+^ release^20^ to activate CaMKII, and the spread of cAMP to activate PKA in nearby synapses. Such a model is consistent with observations that CaMKII and PKA can regulate both functional and structural LTP. It seems most likely that these same mechanisms trigger the heterosynaptic structural alterations.

## METHODS

### Preparation of acute hippocampal slices

Thy1-EGFP mice (JAX strain #007788^30^ on a C57BL/6J genetic background, 10-14 weeks) were anaesthetized with isoflurane and euthanised by decapitation in accordance with the Canadian Council on Animal Care (CCAC) guidelines and AUP approved by the Animal Care Committee (The Centre for Phenogenomics). The brain was then removed and placed in ice cold artificial cerebral spinal fluid (ACSF) based cutting solution (in mM) 124 NaCl, 3 KCl, 26 NaHCO3, 1.25 NaH2PO4, 5 MgSO4, 10 D-Glucose and 1 CaCl2 (bubbled with carbogen, 95% O2 and 5% CO2). The cerebellum and occipital lobes were cut with a scalpel blade. The two hemispheres of the brain were then separated and hippocampi from each hemisphere was removed using a spatula. The hippocampi were then placed on a square of 5% agar and placed on the vibratome stage (Leica, VT1200S). Transverse 400 μm slices were made. Slices were then transferred to an incubation chamber containing modified ACSF (in mM) with 2 MgSO4 and 2 CaCl2 (bubbled with carbogen). Slices were allowed to recover at 32°C for 30 min after dissection and for a minimum of 1 h at 22°C (room temperature) before recordings were made.

### Recording of the field excitatory postsynaptic potential (fEPSP)

Hippocampal slices were perfused at 2 mL/min with the modified oxygenated ACSF at 30°C. A single bipolar electrode and recording electrode was positioned in the stratum radiatum . Schaffer collaterals were stimulated using a constant current stimulator (0.033Hz frequency, pulse width 0.1 ms; STG 4002, MCS, Multichannel systems, Germany) to obtain evoked synaptic responses. Signals were amplified with Multiclamp 700B (Molecular Devices) and digitized with a BNC-2110 (National Instruments) A/D board at a sampling rate of 20 kHz. Throughout the experiments, paired pulses were applied within each pathway (50 ms inter stimulus interval). The initial slope of evoked fEPSPs (V/s) was monitored and analyzed using WinLTP 11 ^31^.

Following a stable baseline of 30 min, LTP was induced using TBS delivered at the same basal stimulus intensity current and pulse width. An episode of TBS comprised 5 bursts at 5 Hz with each burst composed of 5 pulses at 100 Hz (i.e: 25 pulses in total). For spaced TBS (sTBS), three TBS episodes were delivered with an interstimulus interval of 10 min. For the stop stimulation protocol, basal stimulation was paused in between the TBS episodes except for 4 pulses to monitor the level of STP and 30 min following the delivery of the third TBS episode. Representative sample traces are an average of four consecutive responses, collected from typical experiments. Each experiment was conducted from hippocampal slices of different animals. Here, N refers to the number of slices and animals.

### Pharmacological reagents

Compounds were prepared as frozen stock solutions in water (stored at -30°C). These include FM 4-64 (FM / SynaptoRed C2; Biotium Inc.), ADVASEP-7 (Biotium Inc), D-AP5 (Hello Bio Inc, Princeton, NJ, USA), IEM-1460 (IEM; Hello Bio Inc, Princeton, NJ, USA). Drugs were thawed prior to use and diluted into ACSF at least 20 min before their bath application.

### FM4-64 imaging

Presynaptic boutons in acute slices were loaded with FM4-64, using a method previously described ^15,16,18^. In brief, FM4-64 (5 μM) was washed into the slice and loaded through stimulating the Schaffer collateral pathway 100 times at 0.2 Hz. This labels presynaptic terminals. FM4-64 is an amphiphilic dye that binds to the cell membrane. During Schaffer collateral stimulation, vesicular neurotransmitter release occurs, and this is followed by the formation of replacement vesicles manufactured from the labelled external cell membrane. These FM4-64-labelled vesicles are then isolated from the external cell membranes in the presynaptic bouton. To remove residual dye from the external cell membranes, we proceeded with a 10 min ACSF washout followed by a 5 min application of ADVASEP-7 (0.5 mM; Biotium Inc.) for 5 min, followed by a further 30 min wash in ACSF. This restricts FM labelling to only those internalised membranes that form new vesicles. We then followed a destaining protocol that was achieved by delivery of a train of 200 stimuli (0.2Hz) (Fig. 1D). Imaging parameters were kept constant for each batch of FM 4-64 used to ensure accurate P(r) calculations.

To calculate the intensity of the FM dye, we devised a ratiometric measure to ensure accurate calculations of red/green intensity. From our volume image stacks, we examined the axial intensity profile of fluorescence from EGFP and FM dye, only considering synapses whose peak intensity were within 0.5 micron in depth. We experienced bleed-through from the very intense EGFP signal in the postsynaptic spines into the red channel used to detect FM4-64, and therefore ran a linear regression to calculate the bleed through from the red into the green channel and subtracted this to get the red fluorescence signal. P(r) was calculated according to the method described by Murthy et al., 1997^15^ and Sanderson et al., 2018^16^. To calculate the fluorescence intensity that corresponds to a single stained vesicle, we sorted the fluorescence intensity of all FM4-64 puncta into a frequency histogram with bin sizes that are expected to be below the fluorescence intensity of a single synaptic vesicle This resulted in multiple peaks, which are separated by a value that corresponds to the fluorescence of a single synaptic vesicle (x). Therefore, given that we delivered 100 stimulations, the maximum fluorescence that can be achieved is 100x. The P(r) of any given presynaptic puncta is equivalent to the fluorescent intensity off that puncta after 100 stimulations (F) divided by 100x. Therefore, P(r) = F/100x.

### EGFP and FM4-64 imaging

Imaging was performed on a twin, two-photon laser scanning microscope on a FVMPE-RS (Olympus Fluoview, Olympus, Tokyo) equipped with a Mai-Tai Ti:Sapphire Ultrafast Laser (MKS Spectra-Physics) and an Insight X3 laser (MKS Spectra-Physics), and a water immersion objective (×25; 1.05 NA; 2 mm WD; Olympus). Gallium arsenide phosphide photomultiplier tubes (GaAsP PMTs) were used for detection. Images were taken at an additional 6.7× zoom. An excitation wavelength of 920 nm was used. To look at FM staining and destaining, two 512x512 pixel stacks were obtained. The minimum light intensity was used that resulted in adequate signal to noise ratio to prevent photo damage to neurons.

The imaging protocol to study spine morphology during synaptic plasticity consisted of 7 stacks – 2 baseline images, and further images at 0 min, 30 min, 60 min, 90 min and 120 min after TBS. ∼45-50 μm z-stacks were taken at a z-step of 0.5 μm/slice. Z-stacks were taken at ∼20-50μm from the top of the slice with an image size of 1024x1024 pixels to maximize resolution of the images. All images were taken with 2× averaging to reduced noise.

### Image analysis

Fiji/ImageJ^32^ was used to segment and make substacks for each dendrite. To reduce z-drift, NoRMCorre^33^was used to register all the substacks to the first stack. Rigid registration was then performed to remove bulk motion. Subsequently, piecewise rigid registration was performed on the rigid-registered substacks to further co-localize dendritic spines across timepoints.

Registration warps found on the maximum z-projection of the substacks were then applied across all z-levels of the stack. Masks for the dendritic branch as well as dendritic spines were drawn using the cell magic wand plugin on Fiji/ImageJ ^32^ . The centroid of each pixel-wise spine mask was used as the initial search position for each spine. For each timepoint, the previously found mask was used as the initial position to search for the dendritic spine. The peak average pixel intensity across z-levels was used as the expected vertical position for the spine. An average image was created by averaging the pixel intensities ± 1 z-level around the expected z-level for the dendritic spine. A meijering filter was applied to the average-image to create a bias-image around transition points between points of high intensity contrast. The bias-image was then applied to the average-image to negatively bias low-intensity pixels around high-intensity structures. Pixels belonging to the dendrite mask were subtracted out of the biased image to prevent incorrect fitting to the dendrite.

In order to locate FM-positive spines, the spine mask from the green channel was enlarged by 10% and applied to the red channel. If average fluorescence intensity on the red channel was above the noise level within these enlarged ROIs, the spine was denoted as FM+. The centroid of each pixel-wise spine mask in the green channel (EGFP) was used as the initial lateral search position and the peak average pixel intensity in the green channel (EGFP) across z-levels was used as the expected vertical position to locate puncta in the red channel that correspond to connected presynaptic boutons loaded with FM.

A sub-image around the previous position of the dendritic spine was used to fit a 2-D Gaussian and a custom written python script was used for quality-control. All spines were normalised to the average dendrite intensity for each timepoint to compensate for bleaching. For subsequent timepoints when only the EGFP signal was being measured, the previously found mask was used as the initial position to search for the dendritic spine.

### Quantification and statistical analyses

All treatment groups were interleaved with control experiments. Data are presented as mean ± s.e.m. (standard error of mean). In all time-course LTP plots shown, fEPSPs were normalized to the average of the initial 10 min of fEPSP data. Bar plots quantify the synaptic fEPSP data averaged over the final 10 min of recording. In any other instances of fEPSPs averages shown, the time (*t*) in minutes refers to a 10 min bin of data centred over the indicated time. In all time course sLTP plots shown, spine volumes were normalised to the first 2 baselines (0 and 30 min).

Statistical significance was determined using a two-tailed Student’s *t*-test or one-way ANOVA followed by post hoc Dunnett’s multiple comparisons test. Correlation was examined by Pearson’s correlation. All statistical tests were performed using GraphPad Prism (v.10.0.03 for macOS, GraphPad Software, Boston, MA, USA) or MATLAB 2023b. Level of significance within the figures is denoted as follows: \**p* < 0.05; \*\**p* < 0.01; \*\*\**p* < 0.001.

## ACKNOWLEDGEMENTS

This research was funded by CIHR (Canadian Institutes of Health Research) Foundation Grant #154276 to G.L.C. In addition, G.L.C. acknowledges the research infrastructure support from the Canada Foundation for Innovation (CFI) John R. Evans Leaders Fund (JELF, Project #34679) and Ontario Research Fund Program from the Ministry of Research and Innovation (MRI), along with support from the Infrastructure Operating Fund (IOF). G.L.C is the holder of the Krembil Family Chair in Alzheimer’s Research. We thank the Dani Reiss Family Foundation (Neurodegeneration and Aging Research Program) for their support. The work was supported by the Whitehall Foundation and Alfred P. Sloan Foundation (to B.S.)

## AUTHOR CONTRIBUTIONS

L.A.K.: conceptualization, investigation, methodology, formal analysis, visualization, writing–original draft, writing–editing; T.M.S.: conceptualization, supervision, writing–original draft, writing–review and editing; J.G.: conceptualization, supervision, writing–original draft, writing–review and editing, funding acquisition, project oversight. G.B.: image analysis and software tools. B.S. supervision, writing–review and editing, funding acquisition, imaging analysis oversight. G.L.C.: conceptualization, supervision, writing–original draft, writing–review and editing, funding acquisition, project oversight.

